# Accumulation of the GSK3 target protein β-catenin is lethal for B cell precursors and malignant B cells

**DOI:** 10.1101/2021.09.17.460779

**Authors:** Huda Jumaa, Klaus M. Kistner, Corinna Setz, M. Mark Taketo, Hassan Jumaa, Julia Jellusova

## Abstract

Glycogen synthase kinase 3 (GSK3) is a ubiquitously expressed kinase involved in a myriad of biological processes. Although GSK3 mediated phosphorylation has been shown to induce the degradation of many pro-survival and pro-proliferation factors, cancer cells of different origin show reduced proliferation or survival after GSK3 inhibition. Our current understanding of the role GSK3 plays in normal mature B cells, B cell precursors and transformed B cells is incomplete and does not allow to assess whether GSK3 inhibitors can be used to treat B cell derived malignancies. Here we identify β-catenin as the major factor driving GSK3-inhibition induced changes in B cells. We show that β-catenin accumulation has opposing effects on cell metabolism and survival in mature B cells and B cell precursors. Moreover, we demonstrate that β-catenin destabilizes the commitment to the B cell lineage. In summary, our study identifies β-catenin induced signaling as a factor that can be exploited to limit the survival of malignant B cells.

## Introduction

GSK3 is a serine/threonine kinase expressed in α and β isoforms in most mammalian tissues. More than 100 substrates are known to be phosphorylated by GSK3, many of which play important roles in regulating cell metabolism, proliferation and survival(Sutherland, 2011). GSK3 is an atypical kinase in that it is constitutively active in resting cells and disabled upon mitogenic stimulation. In many cell types GSK3 inhibits proliferation by targeting pro-survival and pro-proliferation factors such as cMyc(Gregory et al., 2003) or β-catenin(Doble et al., 2007) for degradation. Despite its role in maintaining cellular quiescence in resting cells, GSK3 has been shown to act both as tumor suppressor and promoter in different types of cancer(Mancinelli et al., 2017). Conflicting reports exist on the role of GSK3 in B cell derived lymphoma. Chemical inhibition or genetic deletion of the GSK3 homologs in different B cell lymphoma lines has been demonstrated to reduce proliferation and to result in cell cycle arrest(Wu et al., 2019). In contrast, B cell receptor mediated GSK3 inhibition has been suggested to increase metabolic fitness in a mouse model of Myc-driven lymphoma. GSK3 inhibition supported lymphoma proliferation and competitive fitness of lymphoma B cells in this model(Varano et al., 2017). It remains unclear, whether the GSK3-signaling context of the various lymphoma cells, the use of different chemical compounds with potential side effects, or the general experimental setup cause the opposing outcomes in these studies. It is conceivable that GSK3 inhibition has various effects on lymphoma cells depending on their specific signaling profile or their maturation and developmental status. Thus in order to harness the full potential of GSK3 inhibitors for B cell lymphoma or leukemia therapy it is imperative to define the conditions under which GSK3 limits B cell survival and proliferation and to clarify the molecular mechanisms driving the phenotype downstream of GSK3.

In normal B cells, GSK3 has been shown to play a dual role in B cell proliferation and survival. GSK3 inhibition in mature activated B cells results in enhanced metabolic activity, increased oxygen consumption, cell mass accumulation and proliferation if nutrients are sufficient(Jellusova et al., 2017). However, the viability of GSK3-deficient B cells is significantly compromised under nutrient restricted conditions in comparison to their normal counterparts(Jellusova et al., 2017). Additionally, the inability to inactivate GSK3 has been demonstrated to result in decreased fitness in response to DNA double strand brakes(Thornton et al., 2016). In conclusion, whether GSK3 supports or limits mature B cell survival and proliferation appears to depend on the context highlighting the multifaceted and dynamic nature of GSK3-dependent signaling.

The role of GSK3 in early B cell development has not been described to this date. Here we analyze how GSK3-inhibition affects B cell precursors, mature B cells and malignant cells and identify β-catenin as a major factor in determining the functional outcome of GSK3 inhibition.

## Results

### GSK3 inhibition reduces proliferation and survival of malignant B cells

To analyze the effect of GSK3 inhibition on B cell derived malignancies, we treated a panel of lymphoma cells of different origin with the GSK3 inhibitor LY2090314. Consistent with previous reports(Wu et al., 2019), GSK3 inhibition resulted in decreased proliferation of lymphoma cells from the lines: Ramos (Burkitt Lymphoma), Jeko (Mantle Cell Lymphoma) and Mino (Mantle Cell Lymphoma) (Fig.1A). OCI-Ly7 (Diffuse Large B cell Lymphoma) and OCI-LY19 (Diffuse Large B cells Lymphoma) cells also showed a slight reduction in proliferation after GSK3 inhibition (Fig.1A). Proliferation of OCI-Ly1 (Diffuse Large B cell Lymphoma) cells was not affected (Fig.1A). While the viability of the tested lymphoma cells was largely not affected after 1 day of culture (Fig.S1A), cell survival was significantly decreased after two days of culture in all cell lines investigated (Fig.1B). To extend our studies to malignant B cells originating from B cell precursors we transformed pre B cells with the oncogenic tyrosine kinase BCR-Abl. This fusion protein can frequently be found in adult acute lymphoblastic leukemia (ALL) patients and is the product of a chromosomal translocation termed Philadelphia chromosome(El Fakih et al., 2018; Pear et al., 1998). We treated the transformed cells with LY2090314 to inhibit GSK3. As expected, GSK3 inhibition in transformed pre B cells resulted in robust accumulation of the GSK3 target β-catenin (Fig.1C). Similar to lymphoma cells the proliferation and survival of transformed pre B cells was decreased and cell death was increased after treatment with LY2090314 (Fig.1D, E).

**Figure 1.:**
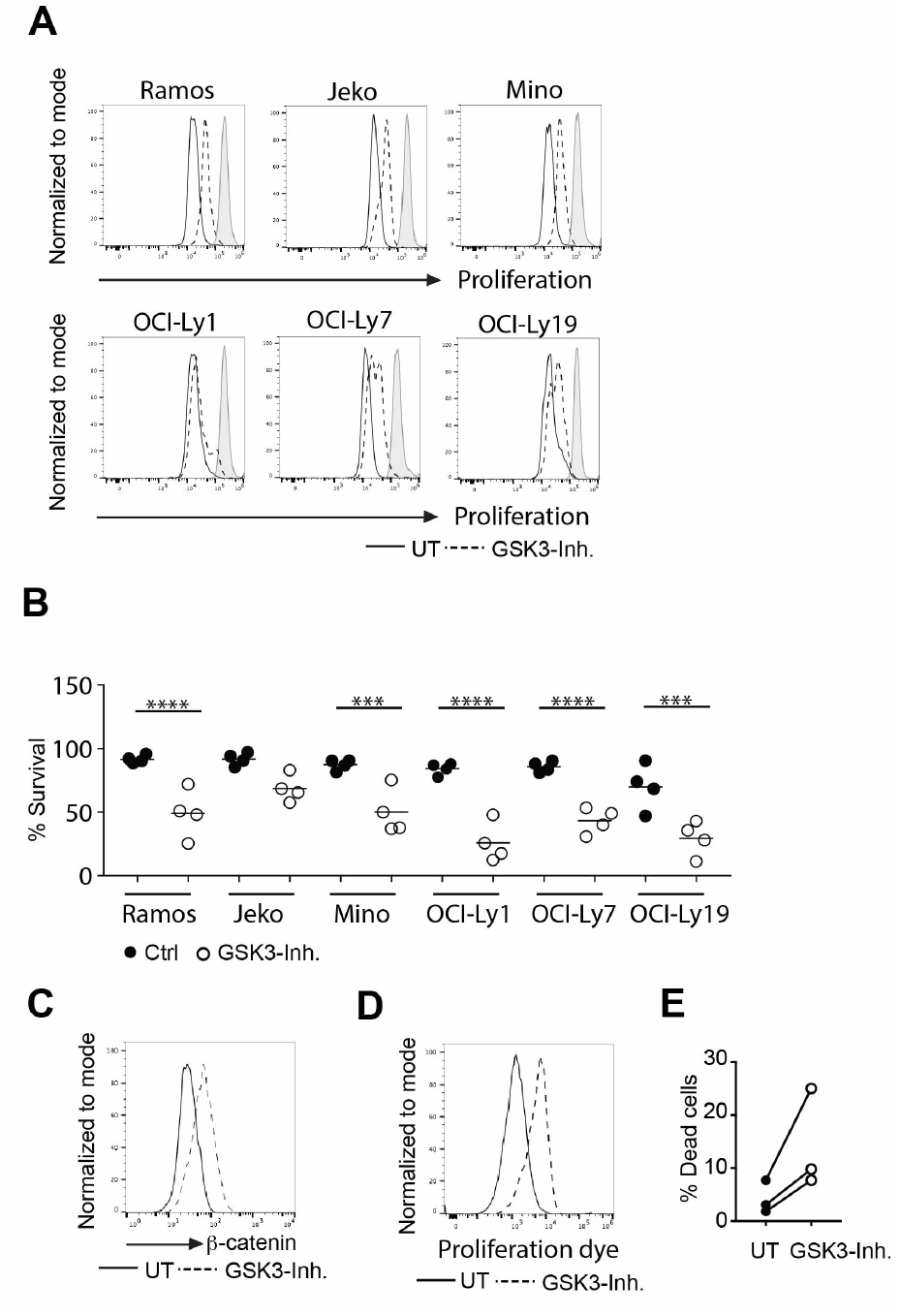
GSK3 inhibition reduces proliferation and survival of malignant B cells. A+B) The indicated lymphoma cell lines were treated with LY2090314 for two days and the dilution of the dye eFluor670 as a measure of proliferation was analyzed by flow cytometry (A). Shown is one out of 3-4 independent experiments. The percentage of viable cells was determined using forward and side scatter. The summary of the performed experiments is shown in B). Significance was determined using the ANOVA test ****p=<0.0001, ***p=0.0002 and 0.0006 C) BCR-Abl transformed B cell precursors were treated with LY2090314 over night. β-catenin expression was analyzed by flow cytometry. One of 3 independent experiments with 5 mice in total is shown. D) BCR-Abl transformed B cell precursors were treated with LY2090314 for 3 days and the dilution of the dye eFluor670 as a measure of proliferation was analyzed by flow cytometry. One of 5 independent experiments is shown. E) Cells were treated with LY2090314 for 3 days. Cell survival was determined by Live/Dead Fixable yellow cell stain incorporation. Circles represent independent experiments. UT= untreated cells GSK3-inh= cells treated with LY2090314

### GSK3 plays a context dependent role in B cells

In previous studies GSK3-deletion has been found to boost the proliferation of mature activated B cells(Jellusova et al., 2017) which is in stark contrast to the phenotype observed in malignant B cells. To rule out general toxicity of LY2090314 and to confirm that GSK3 inhibition causes a similar phenotype as *Gsk3α*/*β* gene deletion we stimulated mature B cells with anti-CD40+IL-4 and treated them with the inhibitor. Since the role of GSK3 in B cell precursors has not been defined before, we included normal, IL-7-dependent B cell precursors in our experiments. Treatment with LY2090314 resulted in robust β-catenin accumulation in both mature stimulated B cells and in B cell precursors (Fig.2A, B). Consistent with the phenotype reported for GSK3-deficient B cells, mature stimulated B cells showed increased proliferation upon GSK3 inhibition (Fig.2C). In contrast, GSK3-inhibition reduced the proliferation of B cell precursors (Fig.2D). GSK3 inhibits mitochondrial activity and reactive oxygen species (ROS) production in mature B cells(Jellusova et al., 2017). To test whether changes in the proliferative rate after GSK3 inhibition are associated with altered mitochondrial activity we measured oxygen consumption and ROS production after GSK3 inhibition. We found basal oxygen consumption, spare respiratory capacity and ROS production to be increased in mature B cells after GSK3 inhibition. However, these parameters were decreased in B cell precursors (Fig.2E, F, G, H). Similar to normal B cell precursors, transformed B cell precursors showed reduced oxygen consumption and ROS production after GSK3-inhibition (Fig.2I, J). Notably, while oxygen consumption was reduced, glucose uptake was increased after GSK3 inhibition (Fig.2K) suggesting that not all metabolic pathways are blocked by GSK3 inhibition. In summary our results show that GSK3 inhibition has different effects on normal mature B cells vs. malignant B cells and B cell precursors.

**Figure 2.:**
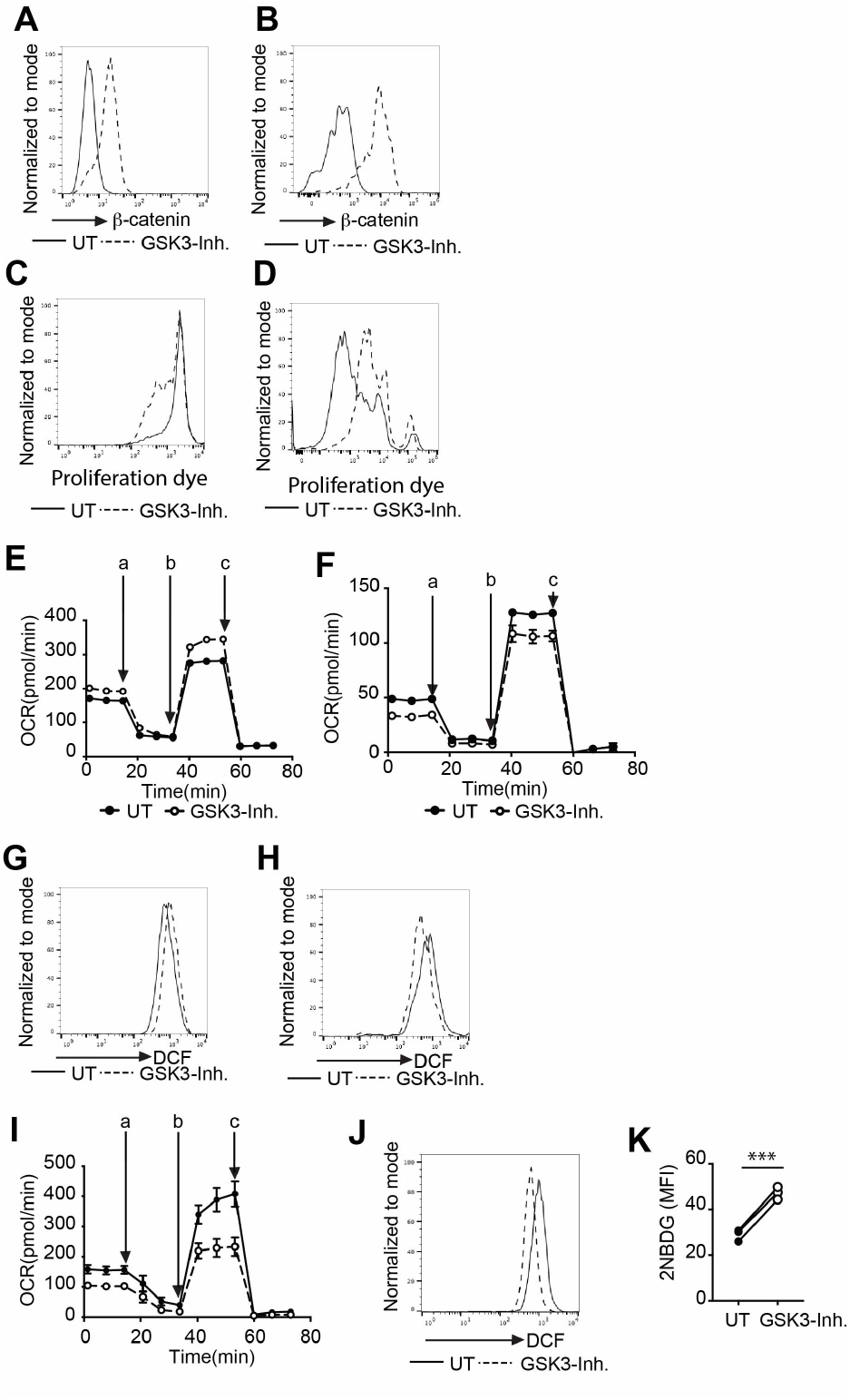
GSK3 plays a context dependent role in B cells. A) Mature B cells were stimulated over night with anti-CD40+IL-4 in the presence or absence of LY2090314, β-catenin accumulation was determined by flow cytometry. One of 6 representative experiments is shown. B) B cell precursors were cultured with IL-7 in the presence or absence of LY2090314 for 1-3 days, β-catenin accumulation was determined by flow cytometry. One of 6 representative experiments is shown. C) Mature B cells were stimulated with anti-CD40+IL-4 in the presence or absence of LY2090314 for 3 days, dilution of the dye eFluor670 as a measure of proliferation was assessed by flow cytometry. One of 6 independent experiments is shown. D) B cell precursors were cultured with IL-7 in the presence or absence of LY2090314 for 3 days, dilution of the dye eFluor670 as a measure of proliferation was assessed by flow cytometry. One of 4 independent experiments is shown. E) Mature B cells were stimulated with anti-CD40+IL-4 in the presence or absence of LY2090314 for 1 day, oxygen consumption was measured. One of 3 independent experiments is shown. F) B cell precursors were cultured with IL-7 in the presence or absence of LY2090314 for 1 day, oxygen consumption was measured. One of three experiments is shown. G) Mature B cells were stimulated with anti-CD40+IL-4 in the presence or absence of LY2090314 for 1-2 days, ROS production was measured by flow cytometry. One of 4 independent experiments is shown. H) B cell precursors were cultured with IL-7 in the presence or absence of LY2090314 for 1-2 days, ROS production was measured by flow cytometry. One of 3 independent experiments is shown. I) BCR-Abl transformed B cell precursors were treated overnight with LY2090314 and oxygen consumption was measured. J) BCR-Abl transformed B cell precursors were treated overnight with LY2090314. ROS production was measured by flow cytometry. One of 3 independent experiments is shown. K) BCR-Abl transformed B cell precursors were treated with LY2090314. Glucose uptake was determined by flow cytometry. Circles represent independent experiments. Significance was determined using the paired t test N=3 p*=0.0008. In experiments in which technical replicates were used circles represent the mean value of these replicates (E, F, I). OCR= oxygen consumption rate, MFI = geometric mean fluorescence intensity, UT= untreated cells GSK3-inh= cells treated with LY2090314, a= oligomycin, b=FCCP, r+a= rotenone + antimycin.

### β –catenin but not cMyc accumulates after GSK3 inhibition in malignant B cells

Reduced mitochondrial activity in GSK3-inhibited malignant B cells and B cell precursors was surprising considering that GSK3 deletion/inhibition has been reported to increase signaling through β-catenin, cMyc and mTORC1 (Doble et al., 2007; Gregory et al., 2003; Inoki et al., 2006) which are central regulators of cell metabolism (El-Sahli et al., 2019; Miller et al., 2012; Saxton and Sabatini, 2017). To identify the molecular mechanisms of reduced mitochondrial activity upon GSK3-inhibition we assessed the protein levels of β-catenin and cMyc as well as S6 phosphorylation, which is a readout of mTORC1 signaling. Similar to what has been reported for GSK3-deficient B cells(Jellusova et al., 2017), mature B cells showed some accumulation of cMyc after GSK3 inhibition and no changes in S6 phosphorylation (Fig.3A). In contrast, we found the levels of cMyc and pS6 to be reduced after GSK3 inhibition in normal and transformed B cell precursors (Fig.3B,C). Similarly, we found β-catenin to accumulate in all cell lines investigated (Fig.3D). cMyc and pS6 levels remained either constant or were reduced after GSK3 inhibition (Fig.3D). In summary our findings demonstrate that the bona fide GSK3 target cMyc does not accumulate in B cell precursors and malignant B cells treated with the GSK3 inhibitor.

**Figure 3.:**
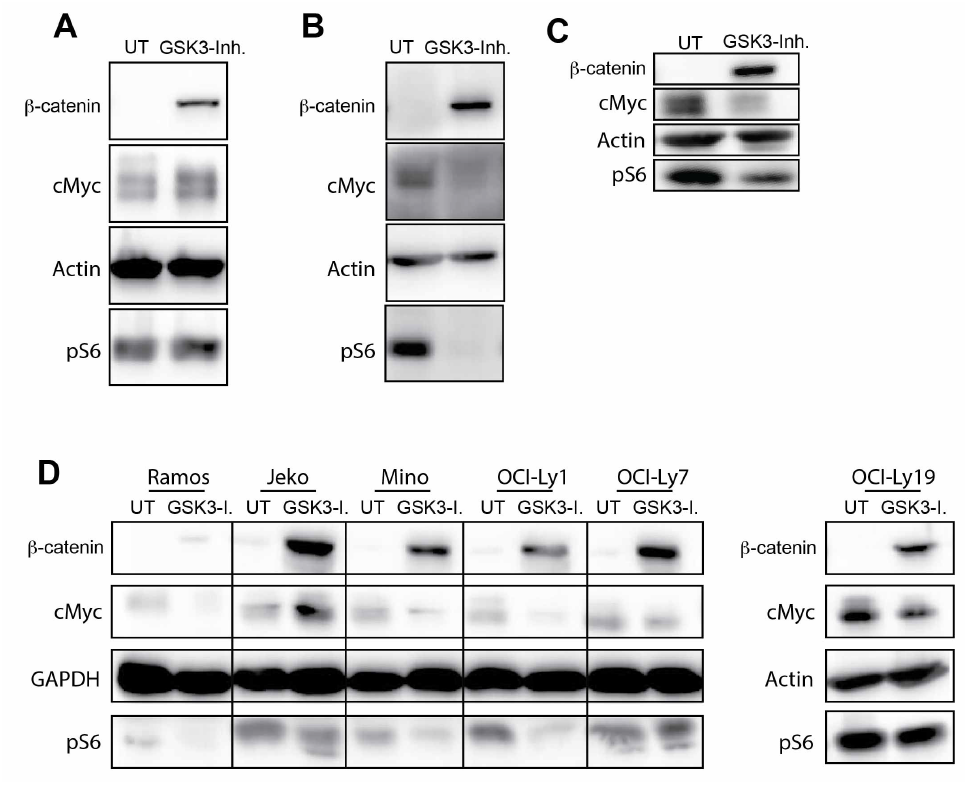
β –catenin but not cMyc accumulates after GSK3 inhibition in malignant B cells. The expression of the indicated proteins was determind by western blot. A) Mature B cells were stimulated with anti-CD40+IL-4 in the presence or absence of LY2090314 for 1 day. One of 3 independent experiments is shown. B) B cell precursors were cultured over night in IL-7 in the presence or absence of LY2090314. One of 5 independent experiments is shown. C) BCR-Abl transformed B cell precursors were treated with LY2090314 over night. One of 2 independent experiments with three mice in total is shown. D) The indicated lymphoma cell lines were treated with LY2090314 over night. One out of three independent experiments is shown. UT= untreated cells GSK3-I= cells treated with LY2090314

### Accumulation of β -catenin interferes with B cell development

Myc is a direct target of GSK3, described to be stabilized after GSK3 inhibition(Gregory et al., 2003), yet failed to accumulate in GSK3 inhibited malignant B cells and B cell precursors. Thus we reasoned that the overall reduction of cMyc proteins levels and S6 phosphorylation are secondary to cell death signals induced by a different GSK3 target molecule. Since we found β-catenin to accumulate consistently in all analyzed B cell subsets we hypothesized that β-catenin plays a central role in orchestrating GSK3-dependent signaling. To analyze the effect of β-catenin accumulation on B cell function, we employed a mouse model in which exon 3 of the β-catenin gene is flanked by loxP alleles and deleted upon Cre enzyme activity (*bCat*^*LoxEx3*^)(Harada et al., 1999). The resulting mutant protein cannot be phosphorylated by GSK3 anymore and accumulates in the cells. To induce β-catenin accumulation in mature B cells, we crossed the *bCat*^*LoxEx3*^ mice to *Mb1-Cre*^*ERT2*^ mice(Hobeika et al., 2015; Hug et al., 2014). *bCat*^*LoxEx3*^ *x Mb1-Cre*^*ERT2*^ mice were injected with tamoxifen to induce Cre activity in B cells. Mature B cells obtained from these mice were stimulated with anti-CD40 and IL-4. Analysis of β-catenin protein levels showed that the majority of B cells accumulated higher levels of β-catenin than control B cells (Fig.4A). Anti-CD40+IL-4 stimulated B cells from *bCat*^*LoxEx3*^ *x Mb1-Cre*^*ERT2*^ mice showed increased proliferation (Fig.4B), oxygen consumption (Fig.4C) and ROS production (Fig.4D) in comparison to cells from control mice, demonstrating that β-catenin accumulation phenocopies GSK3-inhibition induced changes in B cell biology. To analyze the role of β-catenin in B cell development *bCat*^*LoxEx3*^ mice were crossed to *Mb1-Cre* mice(Hobeika et al., 2006). In this mouse model exon 3 of the β-catenin gene is deleted beginning at the pro B cell stage. To analyze B cell development and survival upon β-catenin accumulation, we first assessed the B cell compartment in peripheral lymphoid organs. We found the number of B220-positive cells to be dramatically reduced in the spleen, the lymph nodes and the payers patches (Fig.4E, F and S2A, B, C). Analysis of the expression of maturation markers CD21 and CD23 revealed that few mature B cells (CD21^int^, CD23^high^) reached the periphery (Fig.4G). Similar to peripheral lymphoid organs, mature recirculating B cells (CD43^-^, B220^high^, IgM^+^) were nearly completely absent from the bone marrow of *bCat*^*LoxEx3*^ *x Mb1-Cre* mice (Fig.4H, I). The numbers of small pre B cells (CD43^-^, B220^low^, IgM^-^) and immature B cells (CD43^-^, B220^low^, IgM^+^) were also significantly reduced in *bCat*^*LoxEx3*^ *x Mb1-Cre* mice in comparison to control mice (Fig.4H, I). In contrast, the total cell numbers of pro B cells (CD43^+^, B220^low^, IgM^-^, BP1^-^) and large pre B cells (CD43^+^, B220^low^, IgM^-^, BP1^+^) were slightly but not significantly reduced (Fig.4I, J), suggesting that β-catenin accumulation blocks B cell development at the pre B cell stage.

**Figure 4:**
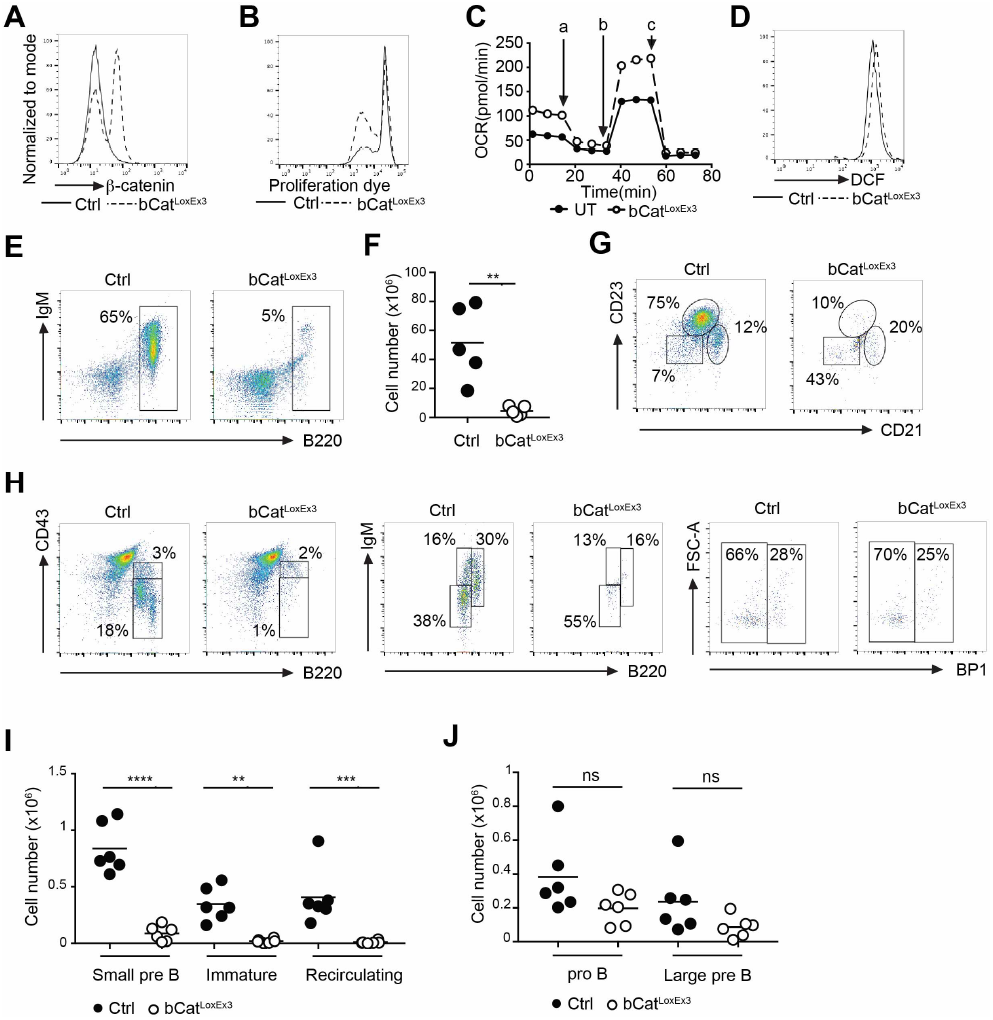
Accumulation of β-catenin interferes with B cell development. A) *Mb1-cre*^*ERT2*^ x *Catnb lox(ex3)* mice were injected 3 times with tamoxifen. Accumulation of β-catenin was determined by flow cytometry. One of at least 6 experiments is shown. B) To induce cre activation *Mb1-cre*^*ERT2*^ x *Catnb lox(ex3)* mice were injected 3 times with tamoxifen or purified B cells were cultured with hydroxy tamoxifen. Mature B cells were cultured with anti-CD40+IL-4 for 3 days. Dilution of the dye eFluor670 as a measure of proliferation was assessed by flow cytometry. One of 6 independent experiments is shown. C) Purified B cells were treated with hydroxytamoxifen and stimulated with anti-CD40+IL-4 for 2 days. Oxygen consumption was measured. One of 3 independent experiments is shown. a=oligomycin, b=FCCP, c= rotenone+antimycin D) *Mb1-cre*^*ERT2*^ x *Catnb lox(ex3)* mice were injected 3 times with tamoxifen. Mature B cells were purified and cultured overnight with anti-CD40+IL-4. ROS production was measured by flow cytometry. One of 3 independent experiments is shown. E+F+G) B cell maturation in spleens from *Mb1-cre* x *Catnb lox(ex3)* mice was analysed by flow cytometry. The frequency of B220^+^ cells was determined. A representative plot is shown in E) total numbers of B220^+^ cells from 5 independent experiments are shown in F). Statistical significance was determined using the unpaired t test, n=5, p** =0.0036 A representative plot showing the frequency of mature B cells (B220^+^, CD21^med^, CD23^+^), marginal zone + transitional type 2 B cells (B220^+^, CD21^+^, CD23^low^) and transitional type 1 B cells + B1a cells (B220^+^, CD21^-^, CD23^-^) is shown in G). One of 5 independent experiments is shown. H+I+J) B cell development in the bone marrow of *Mb1-cre* x *Catnb lox(ex3)* mice was determined by flow cytometry. Representative plots are shown in H), total cell numbers of mature recirculating B cells (CD43^-^, B220^high^, IgM^+^), small pre B cells (CD43^-^, B220^low^, IgM^-^), immature B cells (CD43^-^, B220^low^, IgM^+^), pro B cells (CD43^+^, B220^low^, IgM^-^, BP1^-^) and large pre B cells (CD43^+^, B220^low^, IgM^-^, BP1^+^) obtained from 6 independent experiments are shown in I and J. Statistical significance was determined using the one way ANOVA, n=6, p****<0.0001, p***=0.0002, p**=0.0026 Circles represent independent experiments. Ctrl= control mice, bCatloxEx3 = Cre positive mice carrying the *Catnb lox(ex3)* locus.

### Accumulation of β -catenin induces cell death in B cell precursors

Large pre B cells represent a highly proliferative population of cells and need to become quiescent in order to progress in their development(Herzog et al., 2009). To test whether β-catenin accumulation drives excess proliferation in B cell precursors we made use of an experimental setup in which B cell development from hematopoietic stem cells is induced *in vitro* through the sequential withdrawal of the cytokines Flt3-L and SCF and simultaneous activation with IL-7(Baracho et al., 2014). 6-8 days after Flt3-L and SCF withdrawal, the resulting IL-7 dependent cell culture from control bone marrow cells consisted predominantly of B220^+^CD19^+^ B cell precursors (Fig.5A, B). In contrast, significantly fewer B220+ cells were observed in samples originating from bone marrow obtained from *bCat*^*LoxEx3*^ *x Mb1-Cre* mice (Fig.5A, B). Moreover, while virtually all B220+ cells from control bone marrow co-expressed CD19, on average only 27% of the cells from *bCat*^*LoxEx3*^ *x Mb1-Cre* bone marrow were CD19-positive (Fig.5C). This suggests that B cell development is disrupted after β-catenin accumulation. After analyzing cell cycle distribution of the cultured B cells, we found the frequency of cells in the G1/G0 phase to be significantly increased and the frequency of cells in the S phase to be significantly decreased among B cells from *bCat*^*LoxEx3*^ *x Mb1-Cre* bone marrow (Fig.5D), suggesting that B cells with accumulated β-catenin undergo cellular quiescence or apoptosis. To assess whether β-catenin accumulation decreases cell viability we stained for active caspase 3 and found the frequency of apoptotic cells to be significantly increased in the population of B220+ *bCat*^*LoxEx3*^ *x Mb1-Cre* cells (Fig.5E). In summary, these results demonstrate that unlike in mature B cells, β-catenin in B cell precursors leads to cell cycle arrest and apoptosis.

**Figure 5.:**
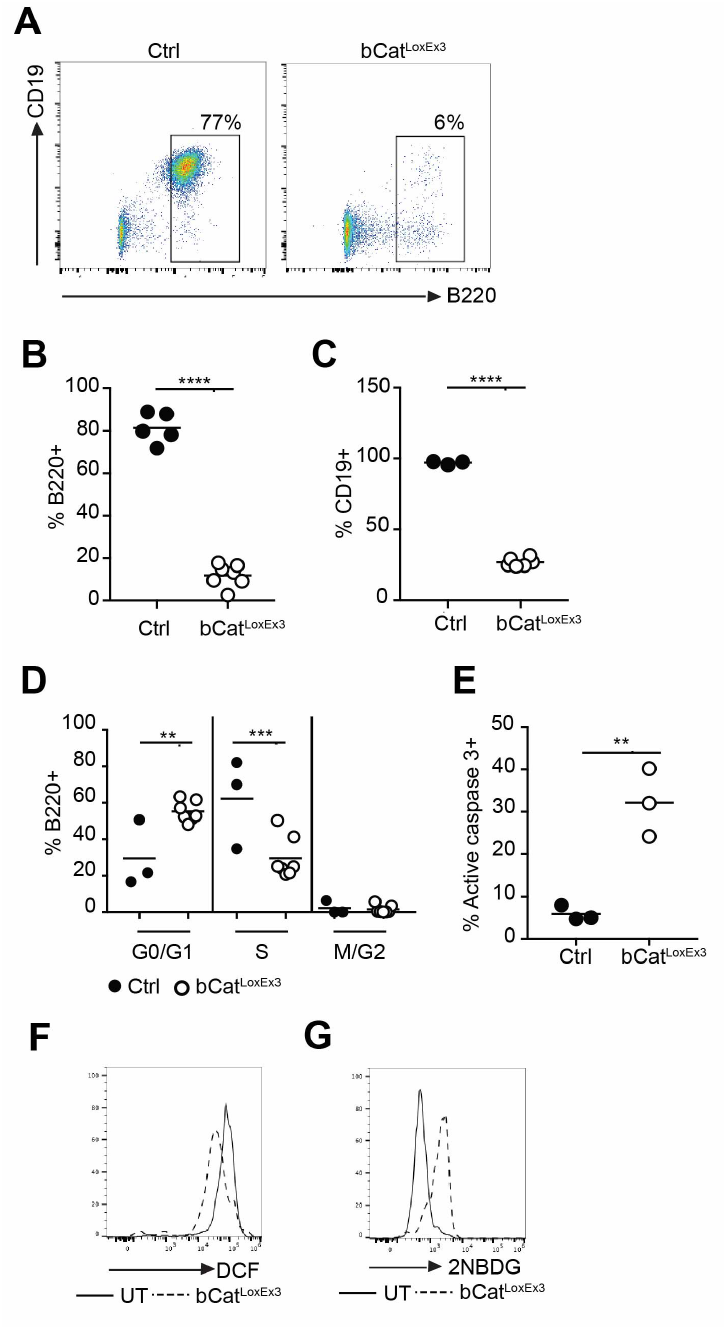
Accumulation of β-catenin induces cell death in B cell precursors. A+B+C) B cell development was analyzed in vitro using *Mb1-cre* x *Catnb lox(ex3)* bone marrow hematopoietic stem cells. The frequency of B220+ cells 7-11 days after the removal of Flt3L+SCF and in the presence of IL-7 was determined by flow cytometry. A representative plot is shown in A). A summary of 5 independent experiments with 5 control mice and 7 experimental mice in total is shown in B). Statistical significance was determined using the unpaired test. N=5 and 7, p****<0.0001 The frequency of CD19^+^ cells within the pool of B220^+^ cells is shown in C). Statistical significance was determined using the unpaired test. N=3 and 6, p****<0.0001 D) Cell cycle distribution in B cell precursors (B220^+^, CD19^+^) obtained from *Mb1-cre* x *Catnb lox(ex3)* bone marrow hematopoietic stem cells cultured in the presence of IL-7 +SCF at day 5-6 of culture was determined by BrdU and 7aad or DAPI incorporation. Statistical significance was determined using the ANOVA test, n=3 and 5, p**=0.0077, p***=0.0008 E) Apoptosis of B cell precursors (B220+, CD19+) obtained from *Mb1-cre* x *Catnb lox(ex3)* bone marrow hematopoietic stem cells cultured in the presence of IL-7 1-5 days after Flt3L+SCF removal was determined by staining for active caspase 3. Statistical significance was determined using the unpaired test, n=3, p**=0.0052 F) ROS production in B cell precursors (B220+, CD19+) obtained from *Mb1-cre* x *Catnb lox(ex3)* bone marrow hematopoietic stem cells cultured in the presence of IL-7 2-8 days after Flt3L+SCF removal was determined using flow cytometry. One of 3 independent experiments is shown. G) Glucose uptake in B cell precursors (B220+, CD19+) obtained from *Mb1-cre* x *Catnb lox(ex3)* bone marrow hematopoietic stem cells cultured in the presence of IL-7 2 days after Flt3L+SCF removal was determined using flow cytometry. One of 4 independent experiments is shown. Circles represent individual mice. Ctrl= control mice, bCatloxEx3 = Cre positive mice carrying the *Catnb lox(ex3)* locus.

Our data demonstrate that similarly to GSK3-inhibited cells, *bCat*^*LoxEx3*^ B cell precursors show reduced proliferation and increased cell death. To further confirm that β-catenin accumulation is a major factor mediating the phenotype observed after GSK3 inhibition, we measured glucose uptake and ROS production in *bCat*^*LoxEx3*^ *x Mb1-Cre* B cell precursors. Similar to transformed B cell precursors (Fig.2J) and normal B cell precursors (Fig.2H) treated with LY2090314, *bCat*^*LoxEx3*^ *x Mb1-Cre* B cell precursors showed reduced ROS production in comparison to control cells (Fig.5F). Furthermore, glucose uptake of *bCat*^*LoxEx3*^ *x Mb1-Cre* pre B cells was increased in comparison to control B cells (Fig.5G). An increase in glucose uptake was also observed in transformed B cell precursors after LY2090314 treatment (Fig.2K). In summary, our data suggest that β-catenin accumulation is a major driving force inhibiting mitochondrial function and survival of B cell precursors after GSK3-inhibition.

### Accumulation of β-catenin disrupts the B cell gene expression profile in B cell precursors

β-catenin can fulfill two separate functions in cells, it can act as a transcriptional co-activator and as a coordinator of cell-cell adhesion(Valenta et al., 2012). To confirm that β-catenin induces changes in the transcriptional profile and to assess which gene clusters are affected by β-catenin stabilization in B cell precursors, we performed transcriptome analysis of B220-positive control and *bCat*^*LoxEx3*^ B cell precursors. Since B220-positive *bCat*^*LoxEx3*^ B cell precursors do not represent a homogeneous population as part of them are CD19 positive and part of them are CD19 negative we also treated wildtype B cell precursors with LY2090314 over night and compared them to untreated samples. We performed principal component analysis (PCA) of the obtained transcriptomes and found that samples obtained from *bCat*^*LoxEx3*^ B cell precursor cultures clustered together and were clearly separated from the control cells (Fig.6A). Similarly, PCA showed a distinct grouping of LY2090314 treated and untreated samples (Fig.6A). The control B cells from the first set of experiments and the untreated samples from the second set of experiments were similar, but not identical in their transcription profile. The control B cells from the first set of experiments were positively sorted for B220 expression, while the untreated cells from the second set of experiments were not sorted prior RNA isolation. These small differences in the experimental setup could possibly explain the observed variations in gene expression profiles between these two sets of samples. The transcription profile of *bCat*^*LoxEx3*^ B cell precursors differed from the transcription profile of LY2090314 treated B cell precursors (Fig.6C), which is expected considering that GSK3 has many other targets beyond β-catenin. However, when we analyzed the set of genes that were significantly upregulated in *bCat*^*LoxEx3*^ B cell precursors when compared to control cells and the set of genes significantly upregulated in LY2090314 treated B cell precursors when compared to untreated cells, we found an overlap of 433 genes (Supplementary table 1 and 2). This set included genes involved in different biological processes such as Wnt signaling and cell differentiation (Supplementary table 3). Of note, we found the expression of *Prdm1*, the gene encoding the differentiation factor Blimp1 to be significantly upregulated in both GSK3-inhibited and *bCat*^*LoxEx3*^ B cell precursors in comparison to their respective controls. Moreover, *Pax5* expression was significantly reduced after GSK3 inhibition. Similarly, there was a trend towards reduced *Pax5* expression in *bCat*^*LoxEx3*^ B cell precursors. PAX5 is a transcription factor defining the B cell lineage (Cobaleda et al., 2007) and Blimp1 is a transcription factor known to drive plasma cell differentiation in mature B cells, but cell death in B cell precursors(Setz et al., 2018). Thus our data indicate that the induction of the GSK3/β-catenin signaling axis destabilizes B cell identity. To verify the results obtained from the transcriptome analysis we first measured Pax5 protein levels in *bCat*^*LoxEx3*^ B cell precursors obtained from hemapoietic stem cell cultures. At day 11 of culture the majority of WT cells expressed the B cell marker B220 and were Pax5 positive (Fig.6B, C). In contrast, the cultures obtained from *bCat*^*LoxEx3*^ hematopoietic stem cells contained only few B220-positive cells (Fig.6B). These cultures also contained a second population of B220-low cells that was not seen in the WT sample. Both B220-low and B220-positive cells from the *bCat*^*LoxEx3*^ culture contained only slightly higher Pax5 levels than B220-negative cells (Fig.6C). These results thus confirm that Pax5 expression is reduced in response to β-catenin accumulation. Moreover both B220-positive and B220-low *bCat*^*LoxEx3*^ cells expressed higher Blimp1 protein levels than B220-positive wildtype cells (Fig.6D). To test whether β-catenin accumulation can disrupt the gene expression profile defining B cell identity in already established B cell precursors, we treated wildtype CD19+ B220+ B cell precursors overnight with LY2090314 and analyzed Pax5 and Blimp1 expression. Strikingly, we found Pax5 expression to be reduced and Blimp1 expression to be increased after GSK3 inhibition (Fig.6E, F). Similar to normal B cell precursors we found Blimp1 expression to increase upon GSK3 inhibition in transformed B cell precursors (Fig.6G) Blimp1 has been reported to induce cell death of B cell precursors(Setz et al., 2018). To test whether Blimp1 is primarily responsible for inducing apoptosis after GSK3 inhibition, we treated control and *Prdm1*-deficient B cell precursors with LY2090314. GSK3 inhibition induced cell death in both WT and *Prdm1*-deficient B cell precursors suggesting that β-catenin has other detrimental effects on cell survival in addition to driving Blimp1 expression (Fig.6H). In our search of additional factors deregulated downstream of β-catenin we found the protein levels of the transcriptional factor Foxo1 to be reduced after GSK3 inhibition in both normal and transformed B cell precursors (Fig.6I, J). Foxo1 is a transcription factor that not only governs essential steps in B cell development but has also been shown to be crucial for the survival and cell cycle progression of acute B cell leukemia(Alkhatib et al., 2012; Dengler et al., 2008; Wang et al., 2018). In conclusion these results demonstrate that β-catenin stabilization not only alters the metabolic profile of B cell precursors but also induces a transcription profile that reverts lineage fate decisions.

**Figure 6.:**
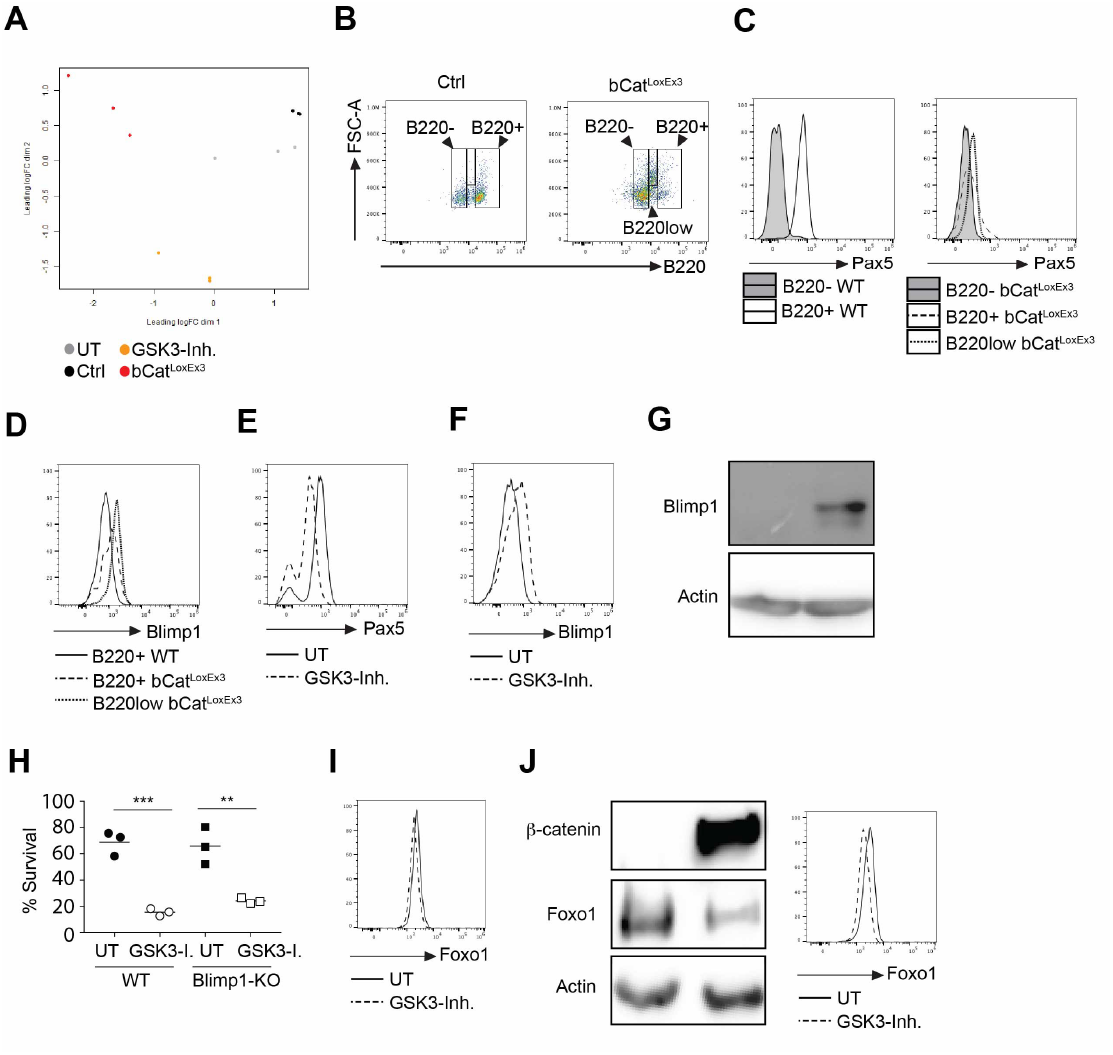
Accumulation of β-catenin disrupts the B cell gene expression profile in B cell precursors. A) PCA analysis of the transcriptome of *Mb1-cre* x *Catnb lox(ex3)* and control B cell precursors and B cell precursors incubated with and without LY2090314 over night. Circles represent samples obtained from different mice. B) Flow cytometric analysis of permeabilized *Mb1-cre* x *Catnb lox(ex3)* and control B cell precursors on day 11 of culture. C+D) Pax5 (C) and Blimp1 (D) expression in B220-, B220+ and B220low cells. Gating strategy shown in B). One out of 3 experiments is shown. E+F) Pax5 (E) and Blimp1 (F) expression in B cell precursors treated with LY2090314. One out of 5 experiments is shown. G) Blimp1 expression in transformed B cell precursors treated with LY2090314. One out of two independent experiments is shown. H) Control and Blimp1-deficient B cells were treated with LY2090314, survival was determined on day 3 via flow cytometry. The ANOVA test was used for statistical analysis **p=0.0016, ***p=0.0003 I) Foxo1 expression in normal B cell precursors treated with LY2090314 as determined by flow cytometry. One of two independent experiments is shown. J) Foxo1 expression in transformed B cell precursors treated with LY2090314 as determined by flow cytometry (left) and westernblot (right). One out of two independent experiments is shown. Circles represent individual mice. Ctrl= control mice, bCatloxEx3 = Cre positive mice carrying the *Catnb lox(ex3)* locus, GSK3-Inh= samples treated with LY2090314, UT= wildtype untreated cells.

## Discussion

B cell derived lymphomas are the most common malignant lymphoid neoplasms (Küppers, 2005; Young and Staudt, 2013) and new strategies are required to treat refractory disease. GSK3 has emerged as a potential new target for medical intervention in different types of cancer. Owing to the complexity of the signaling processes governed by GSK3 it remains however difficult to predict the biological outcome of GSK3 inhibition in B cell derived malignancies. Inactivation of genes encoding both of the GSK3 isoforms results in enhanced oxygen consumption, cell mass accumulation and proliferation in mature stimulated B cells(Jellusova et al., 2017). This phenotype is consistent with the accumulation of the transcription factors cMyc and β-catenin in these cells. Both of these transcription regulators have been shown before to support a transcription profile favoring increased metabolic activity(Dang, 2013; Sherwood, 2015). Similar to mature stimulated cells, B cell lymphoma cells have been shown to benefit from GSK3 inhibition(Varano et al., 2017). Signals originating from the B cell receptor have been suggested to result in GSK3 inhibition, which in turn increases the competitive fitness of the cells(Varano et al., 2017). These findings would thus discourage the use of GSK3 inhibitors for B lymphoma treatment. However, other studies have found GSK3 inhibition to successfully inhibit lymphoma B cell proliferation(Wu et al., 2019). In our study we have confirmed that GSK3 inhibition reduces proliferation and survival in lymphoma cells. The role of GSK3 in transformed B cell precursors has not been studied before and our results demonstrate for the first time that GSK3 inhibition reduces metabolic activity and proliferation of transformed B cell precursors. Thus we have identified a possible new target that could be exploited for therapy in of B cell precursor acute lymphoblastic leukemia.

Moreover, we show that while GSK3 inhibition induces increased mitochondrial activity and faster proliferation in mature stimulated B cells, this treatment exerts the opposing effect on B cell precursors arguing for a context dependent role of GSK3 in B cells. GSK3 possesses a plethora of potential targets and teasing apart the role of these various proteins is essential in order to be able to identify patients that could benefit from GSK3 inhibition. GSK3 has been shown to bind to centrosomes and suggested to play an important role in cell cycle progression(Wu et al., 2019). While it is possible that GSK3 plays a role in mitosis progression, this function is not absolutely necessary for B cells to proliferate as both normal and lymphoma B cells have been shown to be able to undergo cell division in the presence of GSK3 inhibitors or in the absence of the GSK3α/β proteins(Jellusova et al., 2017; Varano et al., 2017). In this study we propose that the outcome of β-catenin activity may be the determining factor of whether GSK3 inhibition results in accelerated proliferation or cell death. Despite the wide array of proteins being dysregulated after GSK3 inhibition, we found that β-catenin accumulation alone is sufficient to replicate the phenotype of GSK3-inhibited cells. Similar to B cells treated with a GSK3 inhibitor, mature B cells with hyper-stabilized β-catenin show increased oxygen consumption, ROS production and proliferation. Equally, both GSK3-inhibited B cell precursors and B cell precursors with hyper-stabilized β-catenin display reduced production of ROS and increased cell death. Of note, we found cMyc protein levels and the phosphorylation of ribosomal protein S6 to be differentially regulated in mature B cells and B cell precursors upon GSK3 inhibition. GSK3 has been previously reported to negatively impact on both cMyc levels (Gregory et al., 2003) and mTORC1 activity (Inoki et al., 2006). In other cell types, GSK3 has been shown to phosphorylate and activate the TSC complex, which in turn inhibits mTORC1. However in B cells, this signaling pathway does not seem to play a dominant role in regulating S6 phosphorylation, since both GSK3-deficient and GSK3-inhibited mature B cells show normal S6 phosphorylation(Jellusova et al., 2017). cMyc is targeted for degradation via GSK3 induced phosphorylation(Gregory et al., 2003) and GSK3 deletion or inhibition has been shown to result in cMyc accumulation in various cell types, consistent with what we have observed in mature B cells. Surprisingly, we have found cMyc levels and S6 phosphorylation to be strongly decreased in GSK3-inhibited B cell precursors and malignant B cells. Since both cMyc and mTORC1 signaling support mitochondrial biogenesis and function, reduced activity of these signaling pathways could explain the observed defects in mitochondrial activity.

In summary our findings suggest that β-catenin plays a dominant role in determining B cell fate and that unlike in many other cell types β-catenin accumulation in B cell precursors dampens metabolic activity and proliferation.

β-Catenin is a transcriptional co-activator and we could show that both GSK3 inhibition or β-catenin accumulation not only alter the metabolic program of B cell precursors, but also dramatically reshape their gene expression profile. Particularly, GSK3 inhibition/β-catenin accumulation resulted in reduced Pax5, Foxo1 and increased Blimp1 expression. Moreover we show that B cell precursors which already demonstrate commitment to the B cell lineage by expressing B220 and CD19 still downregulate Pax5 upon GSK3 inhibition. Blimp1 is a transcription factor that is known to induce B cell differentiation to plasma cells and has recently been demonstrated to negatively impact the survival of B cell precursors(Hug et al., 2014; Setz et al., 2018). Thus, increased Blimp1 expression could contribute to reduced survival of B cell precursors after GSK3 inhibition/β-catenin stabilization. However, since Blimp1 deletion failed to rescue survival of B cell precursors after GSK3 inhibition, other signaling events likely contribute to the observed phenotype. In addition to reduced Pax5 levels, we have observed the protein levels of the transcription factor Foxo1 to be reduced upon GSK3 inhibition. Consistent with the essential role of Foxo1 in B cell development and B-ALL survival (Alkhatib et al., 2012; Dengler et al., 2008; Wang et al., 2018) the observed reduction of Foxo1 protein levels could contribute to impaired survival after GSK3 inhibition.

As a transcriptional activator, β-catenin can pair with different factors including the transcription factors from the TCF/LEF family or Hif1α(Kaidi et al., 2007; Valenta et al., 2012). It is possible that β-catenin interacts with different transcription factors depending on the maturation stage of B cells. Identification of the main β-catenin interaction partners in mature B cells and B cell precursors could thus help to understand the different behavior of these two subsets. Alternatively, β-catenin accumulation could initially drive a hyper-metabolic phenotype supporting proliferation in mature B cells, but quickly deplete cellular energy stores leading to a metabolic collapse in B cell precursors. B cell precursors have been suggested to be particularly sensitive to energetic stress (Müschen, 2019). A model has been proposed in which aberrantly increased metabolic activity is interpreted as a sign of malignant transformation or the expression of an autoreactive BCR and therefore stringent mechanisms must exist to eliminate B cell precursors with high energetic demands(Müschen, 2019). Consistent with this model metabolic stress caused by β-catenin accumulation may signal oncogene activation in pre B cells and thus lead to cell death. In summary, our study provides new insight into GSK3 driven B cell signaling and provides a rationale to focus on β-catenin induced transcriptional changes in order to fully harness the potential of GSK3 inhibitors for the treatment of B cell derived malignancies.

## Material and methods

### Mice

Mice bearing the *Catnb lox(ex3)* locus have been described previously(Harada et al., 1999). *Catnb lox(ex3)* mice were crossed to *mb1-cre*^*ERT2*^ mice (Hobeika et al., 2015; Hug et al., 2014). Mice were i.p. injected with 1mg tamoxifen (Sigma) + 10% ethanol (Roth) in olive oil on three consecutive days. Control animals were injected with tamoxifen the same way as experimental animals. To induce deletion of *catnb* exon 3 in B cell precursors *Catnb lox(ex3)* mice were crossed to *Mb1-cre* mice. For both lines, mice homozygous or heterozygous for the *Catnb lox(ex3)* locus were used as experimental animals. Since no significant differences were observed between homozygous and heterozygous mice, the exact genotype is not indicated when presenting data. As controls, both Cre-positive and negative animals were used. No significant differences were observed between these two types of control animals. Both male and female mice were used for experiments. Animals were maintained in a specific pathogen free environment. Experiments were approved by the regional council in Freiburg, Germany and carried out in accordance with the German Animal Welfare Act. For experiments with Blimp1-deficient B cell precursors, bone marrow was obtained from *Mb1-cre x Prdm1*^*lox*^ mice (Setz et al., 2018).

### B cell purification and cell culture

To obtain mature B cells, spleens were homogenized and red blood cells were lysed using an ammonium-chloride-potassium solution (150mM NH4Cl, 1mM KHCO3, 0.1mM Na2EDTA). B cells were purified using CD43 magnetic beads (Miltenyi) or the EasySep mouse B cell isolation kit (StemCell) following manufacturers instructions. Cells were cultured in RPMI (Gibco) + 10% FBS (Biochrome AG LOT 0340A) + 100Units/ml Penicillin + 100µg/ml Streptomycin (Gibco) + 1 mM sodium pyruvate (Thermo Fisher Scientific) + 2 mM Glutamax + 1x non-essential amino acids (Gibco) + 57µM β-Mercaptoethanol (Sigma) at 37°C in an atmosphere with 5% CO2. The following reagents were used in individual experiments: 5µg/ml anti-CD40 (SantaCruz or BioLegend), 10ng/ml IL-4 (Sigma), 40nM LY2090314 (Sigma).

Two different protocols were used to obtain B cell precursors. In the first experimental setup bones were flushed with 10%FBS (PAN, Lot: P140508) PBS, red blood cells were lysed and cells were cultured in Iscove’s medium (Merck, Biochrom)+10% FBS (PAN Premium Lot: P140508) + 100Units/ml Penicillin + 100µg/ml Streptomycin (Gibco)+2mM Glutamax + 57µM β-Mercaptoethanol (Sigma) + 0.4ng/ml IL-7 (Sigma)+40µl/ml IL-7 (obtained from IL-7 producing J558L cells) at 37°C in an atmosphere with 7.5% CO2. Cells were repeatedly re-plated on new cell culture dishes to remove adherent cells. Experiments were performed once the culture reached a purity of 90-95% CD19+B cells. Cells were kept in culture for up to 8 weeks. This protocol was used to generate B cell precursors used in experiments presented in figures: 2F, 2H and 3B. In the second experimental setup, lineage negative cells were obtained from bone marrow using anti-Gr1, CD11b, CD3e, CD49b, Ter119 and B220 antibodies coupled to biotin (eBioscience) and anti-biotin magnetic beads (Miltenyi). Lineage-depleted cells were cultured in 10ng/ml IL-7 (Sigma), 50ng/ml Flt3-L (Sigma) and 50ng/ml SCF (Sigma) in Opti-MEM medium (Gibco) with: 20%FBS premium (PAN, Lot: P140508)+ 1 mM sodium pyruvate (Thermo Fisher Scientific)+ 2 mM Glutamax +25mM HEPES + 100Units/ml Penicillin + 100µg/ml Streptomycin (Gibco)+ 57µM β-Mercaptoethanol (Sigma) at 37°C in an atmosphere with 7.5% CO2.

Flt3-L was withdrawn after 3 days and SCF was withdrawn after 6-8 days. The cells were cultured in IL-7 alone or IL7+SCF for at least 2 days before experiments were performed. B cell precursors obtained through this protocol were used in experiments shown in figures: 5 and 6. In experiments where GSK3 was inhibited, cells were plated in a concentration of 10^6^/500µl and were treated with 40nM LY2090314 (Sigma). To transform B cell precursors, the cells were transformed retrovirally with *BCR-ABL1* (Pear et al., 1998). IL-7 was removed from the culture to eliminate non-transduced cells. Transformed B cell precursors were maintained in Iscove’s medium (Merck, Biochrom)+10% FBS premium (PAN Lot: P140508)+ 100Units/ml Penicillin + 100µg/ml Streptomycin (Gibco)+2mM Glutamax (with Iscove’s medium without stable Glutamine)+ 57µM β-Mercaptoethanol (Sigma) at 37°C in an atmosphere with 7.5% CO2. To inhibit GSK3, cells were plated in a concentration of 10^6^/ml with or without 40nM LY2090314 (Sigma).

All lymphoma cell lines were maintained in RPMI + 10% FBS + 100Units/ml Penicillin + 100µg/ml Streptomycin (Gibco) + 57µM β-Mercaptoethanol at 37°C in a 5% CO2 atmosphere. To inhibit GSK3, cells were plated in a concentration of 0.5 10^6^/1ml with or without 40nM LY2090314 (Sigma).

### Flow cytometry

Single cell suspensions were stained with fluorescently labeled antibodies in FACS buffer (PBS+ 1% BSA+ 0.09% NaN3). The following antibodies were used for flow cytometry: B220(RA3-6B2, Biolegend or ThermoFisher), BP1(6C3, eBioscience), CD19(eBio1D3, ThermoFisher), CD21/CD35 (4E3, ThermoFisher), CD23(B3B4, eBioscience)and IgM (II/41, eB121-15F9,eBioscience). To distinguish between life and dead cells 7AAD (Sigma) or LIVE/DEAD fixable yellow death stain kit (Molecular Probes) were used. For intracellular staining cells were fixed and permeabilized with 2% paraformaldehyde and 70% methanol or with BD Cytofix/ Cytoperm buffer (BD Biosciences) and permeabilization buffer (eBioscience) and incubated with 5µl β-catenin in 100µl Perm buffer (BD), 0.5µl cleaved caspase 3 (Cell Signaling Technology) or anti-Foxo1 (Cell Signaling Technology) in 100µl Perm buffer (BD) for 1h hour on ice. Cells were washed twice with Perm buffer (BD) and incubated with an anti-rabbit secondary antibody (Biolegend) if needed. For Blimp1 and PAX5 intracellular staining, cells were fixed and permeabilized with 3.1% paraformaldehyde and 0.05% Triton in PBS with 1µl anti-Blimp1 (Biolegend) or 1µl anti-Pax5 (Biolegend) in 100µl Perm/wash buffer (BD) for 1h hour at RT. For cell cycle analysis, 10µM BrdU (eBioscience) was added to the cell culture for 24h. Cells were fixed and permeabilized using the BD Cytofix/Cytoperm buffer system (BD) and permeabilization buffer (eBioscience), incubated with 0.3mg/ml DNAse (eBioscience) for 1h at 37°C, washed with Perm buffer (BD) and treated with anti-BrDU (eBioscience) in Perm buffer for 30min at room temperature. Cells were washed, 0.5µg/ml DAPI or 7AAD was added to detect DNA, and live cells were identified gating on forward side scatter. Cell cycle analysis was performed on day 5-6 of cell cuture, cells were SCF+IL7 dependent at this point.

Cells were analyzed by flow cytometry. The following cytometers were used for acquisition of flow cytometry data: LSR II (BD), CyAn (Beckman Coulter), Attune (Thermo Fisher Scientific). FlowJo software (TreeStar) was used for analysis.

### Analysis of metabolic parameters and proliferation

Oxygen consumption was assessed using a Seahorse XFe96 metabolite analyzer (Agilent). 10^5^ Ramos cells, 1×10^5^ transformed or 3×10^5^ normal B cell precursors or 10^6^ mature stimulated B cells were plated on Cell-Tak (Corning) coated Seahorse cell culture plates. The cells were first incubated in Seahorse base medium + 1mM sodium pyruvate (Thermo Fisher Scientific) + 2mM L-glutamine (Thermo Fisher Scientifc) + 10mM glucose (Sigma) in a volume of 50µl for 30min, 130µl medium were added in a second step and the cells were incubated for an additional 1h. Oligomycin, FCCP and rotenone+antimycin were sequentially injected during the measurement to a final concentration of 1µM each to assess different parameters of respiration. To measure the production of reactive oxygen species, cells were stained with 10µM carboxy-H2DCFDA (Thermo Fisher Scientific) for 20min at 37°C and washed before measurement. To assess glucose uptake cells were incubated with 30µM 2NBDG (Cayman Chemical) in PBS (Invitrogen) for 30min at 37°C and analyzed by flow cytometry. To analyze proliferation, cells were loaded with 5µM proliferation dye eFluor670 (eBioscience) and cultured for up to 3 days. Dilution of the proliferation dye was measured by flow cytometry.

### Immunoblot analysis

Cells were lysed in RIPA buffer (150 mM NaCl, 1% NP-40, 0.5% sodium deoxycholate, 0.1% sodium dodecyl sulfate, 50 mM Tris, 1 mM EDTA) supplemented with proteinase and phosphatase activity inhibitors sodium orthovanadate (1mM), sodium fluoride (10mM) and proteinase inhibitor cocktail containing AEBSF, aprotinin, bestatin hydrochloride, E-64, EDTA, leupeptin (Sigma). Immunoblotting was performed following standard procedures. PVDF membranes (Merck Milipore) were blocked with 5% milk powder in TBS buffer (150 mM NaCl, 50 mM Trizma, pH 7.6)+ 0.1% Tween20. All primary antibodies were purchased from Cell Signaling Technology and used at a 1:1000 dilution. The following antibodies were used: Actin (13E5), β-catenin (D10A8), cMyc (D84C12), pS6^Ser(235/236)^ (D57.2.2E), Blimp1 (C14A4), Foxo1 (C29H4). A horseradish-peroxidase-coupled goat anti-rabbit IgG (Jackson or Cell Signaling Technology) was used as a secondary antibody.

### Transcriptome analysis

B cell precursors were obtained from hematopoietic stem cell cultures of control or *Mb1*^*cre*^x *Catnb lox(ex3)* mice as described above and sorted based on B220 expression by flow cytometry, after sorting cells were directly processed for RNA isolation. For experiments in which GSK3 was inhibited, B cell precursors were treated with LY2090314 over night and directly processed for RNA isolation without prior sorting. RNA was extracted using Quick-RNA MicroPrep (ZYMO RESEARCH) according to the manufacturer’s instructions. The RNA concentration in RNAse free H2O was analyzed by measuring the absorption at 260 / 280 nm with a NanoDrop (Thermo Scientific/Peqlab). Additionally, unsorted wildtype B cell precursors were incubated with or without 40nM LY2090314 (Sigma) over night and RNA was isolated. RNA was snap frozen in liquid nitrogen, samples were processed and the transcriptome was analyzed by Novogene company.

### Statistical analysis

GraphPad Prism was employed for statistical analysis. To test for normal distribution the Shapiro-Wilk normality test was used. For independent normally distributed data the unpaired t test was used. For linked normally distributed data the paired t test was used. Otherwise the Mann Whitney U test was used. The specific test used to evaluate statistical significance is listed in the respective figure legend. Differences were considered statistically significant if *p<0.05. Number of independent repeats and technical replicates are indicated in the figure legend. In all graphs showing statistical analysis the mean is indicated by a small horizontal line. Statistical analysis of transcription data was performed by Novogene company. Geneontology enrichment analysis was performed using PANTHER on the geneontology.org website (Ashburner et al., 2000; Carbon et al., 2019; Mi et al., 2019), principal component analysis was performed and visualized in a multidimensional scaling plot using R statistical software (Chen et al., 2016; R-core-team, 2020). Mice were allocated to groups based on their genotype. Numbers rather than the genotype were used to label samples, however the researchers were not entirely blinded to group allocation. Sample sizes were determined based on previous experience. For experiments shown in figure 4I and 4J a power analysis was performed using the following parameters: effective size=1.7, power=0.8, sig.level=0.05. The effective size was determined based on previous experiments in which large differences were expected to be observed. Using these parameters the n was determined to be 6.54 for each group.

## Data availability

Raw data obtained from transcriptome analysis were deposited with GEO and will be made publicly accessible upon manuscript acceptance. Original, uncropped pictures of shown western blots are included in source data. Additional repeats of the experiments are also included in the source data. Numerical data used for all the shown graphs are included in the source data.

## Acknowledgements

We would like to thank John Apgar, Sanford Burnham Prebys Medical Discovery Institute, La Jolla, USA for providing us with the lymphoma cell lines: Jeko, Mino, OCI-LY1, OCI-LY7 and OCI-LY19. We would like to thank Michael Reth for providing us with the *mb1-cre*^*ERT2*^ and the *Mb1-Cre* mice. We would like to acknowledge the Signalling Factory of the Research Centres BIOSS and CIBSS for providing support with flow cytometry.

## Author contribution

Hu.J performed the majority of the experiments and analyzed the data and contributed to writing the manuscript. K.M.K performed part of the experiments. C.S. and Ha.J. provided bone marrow from *Mb1-cre x Prdm1*^*lox*^ mice. Additionally, Ha.J. contributed to editing the manuscript and provided ideas for experiments. M.M.T provided the *Catnb lox(ex3)* mice. J.J conceived of and coordinated the study, performed part of the experiments, interpreted the results and wrote the manuscript.

---

**Figure S1.:**
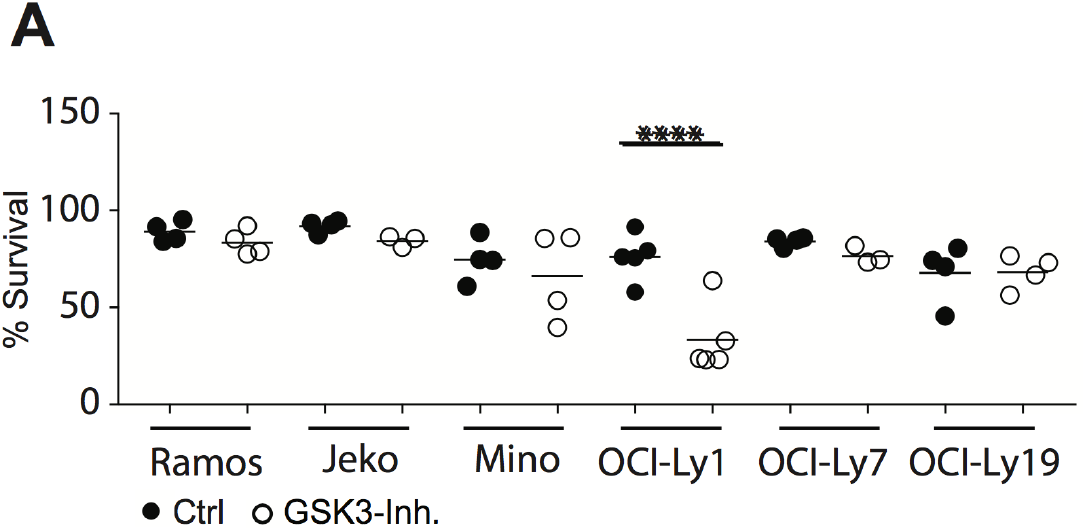
Survival of lymphoma cells after GSK3 inhibition. A) The indicated lymphoma cell lines were treated with LY2090314 for one day. The percentage of viable cells was determined using forward and side scatter. Circles represent independent experiments. UT= untreated cells GSK3-I= cells treated with LY2090314.

**Figure S2.:**
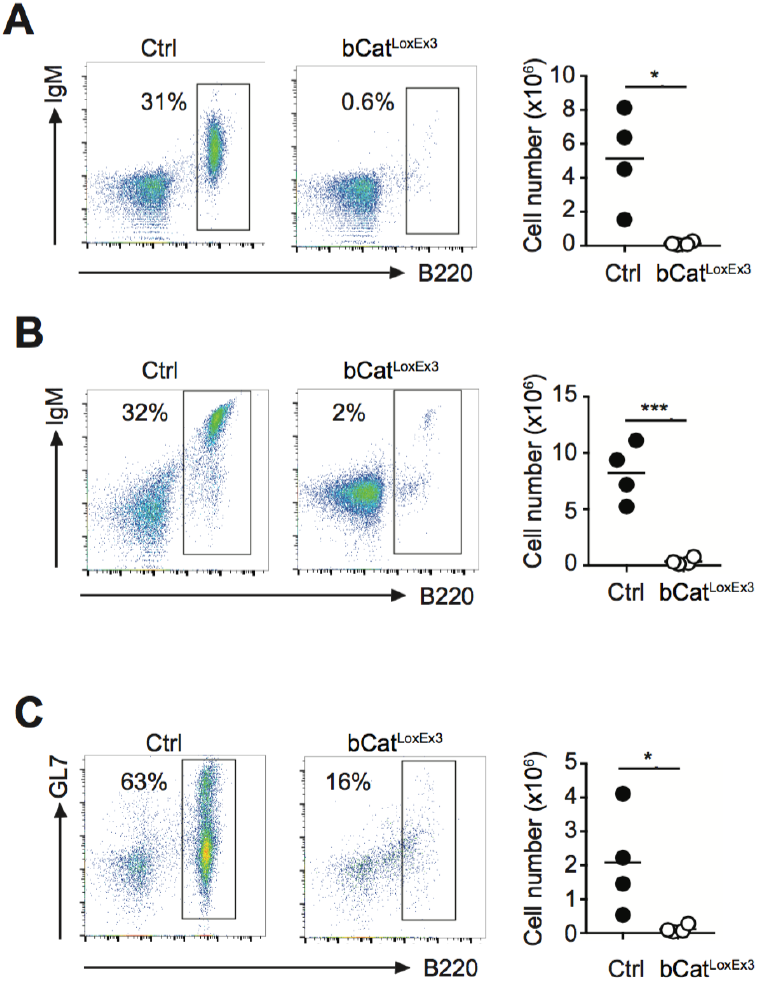
Accumulation of β-catenin results in a loss of mature B cells in the periphery. The frequency of B cells (B220+) in lymphoid organs from *Mb1-cre* x *Catnb lox(ex3)* mice was analyzed by flow cytometry. Shown are representative plots (left) and the summary of absolute B cell numbers obtained in 4 independent experiments (right) from peripheral lymphnodes A) mesenteric lymph nodes B) and Payer’s patches (C). Statistical significance was determined using the unpaired t test. *p=0.012, ***p=0.0009, *p=0.0421 Circles represent independent experiments. Ctrl= control mice, bCatloxEx3 = Cre positive mice carrying the *Catnb lox(ex3)* locus.

## Supplementary Table legend

**Table S1.: Accumulation of β-catenin changes the transcription profile of B cell precursors**

The transcription profile of *Mb1-cre* x *Catnb lox(ex3)* and control B cell precursors and the transcription profile of normal B cell precursors cultured with or without LY2090314 over night was analyzed. The table lists genes for which transcription was significantly increased in *Mb1-cre* x *Catnb lox(ex3)* B cell precursors in comparison to control B cell precursors and genes significantly increased in LY2090314 treated B cell precursors in comparison to untreated B cell precursors.

**Table S2.: Genes overexpressed in both GSK3-inhibited and Catnb lox(ex3)B cell precursors**

The transcription profile of *Mb1-cre* x *Catnb lox(ex3)* and control B cell precursors and the transcription profile of normal B cell precursors cultured with or without LY2090314 over night was analyzed. The table lists genes for which transcription was significantly increased in both *Mb1-cre* x *Catnb lox(ex3)* B cell precursors in comparison to control B cell precursors and in LY2090314 treated B cell precursors in comparison to untreated B cell precursors.

**Table S3.: Genes overexpressed in both GSK3-inhibited and Catnb lox(ex3) B cell precursors are involved in various biological processes**

The transcription profile of *Mb1-cre* x *Catnb lox(ex3)* and control B cell precursors and the transcription profile of normal B cell precursors cultured with or without LY2090314 over night was analyzed. To give a broad overview of biological processes the genes overexpressed in both *Mb1-cre* x *Catnb lox(ex3)* and GSK3 inhibited cells are involved in, we performed a PANTHER overrepresentation analysis. The table lists biological processes in which the overexpressed genes are involved.

## Sources data

The sources data include numerical values underlying graphs shown in Fig.1, Fig.1S, Fig.2K, Fig.4 Fig.5, Fig6H and Fig.2S. This file is labeled “Fig1_2_4_5_6-source data”. The source data also include the original uncropped western blot pictures. The pictures are labeled with the number of the figure and the antibody that was used in the last step. For membranes that have been cut and for which the picture shows several parts incubated with different antibodies, the name of the figure reflects the antibody relevant for the paper. In addition, pdf files are included showing western blots as presented in the figures together with uncut versions indicating the relevant lines with an asterisk. These files also include additional repeats of the representative examples shown in the manuscript. The western blots shown in the manuscript are labeled with the number and letter of the respective figure legend, the repeats are labeled as “repeat”.

